# Angiogenic PolyHIPE Scaffolds Decorated with Extracellular Matrix as Periosteum Substitutes

**DOI:** 10.1101/2025.08.17.670721

**Authors:** Caitlin S. Ryan, Frederik Claeyssens, Gwendolen Reilly

## Abstract

The periosteum is the connective tissue that envelopes bone and contributes to the normal bone healing process. Periosteal grafts have shown excellent success in the treatment of nonunion bone defects but surgical challenges such as donor site morbidity and graft availability have limited their clinical use. Artificial periosteal membranes are being explored as off-the-shelf alternatives, with several groups investigating the use of decellularized tissues or extracellular matrix (ECM)-based strategies for this purpose. In this study, we investigate the use of *in vitro*-generated, fibroblast-derived ECM to decorate porous biodegradable scaffolds for use as synthetic periosteal grafts. Scaffolds were fabricated using an emulsion templating technique from a blend of polycaprolactone methacrylate (PCL-M) and poly(glycerol sebacate) methacrylate (PGS-M), resulting in membranes with large, interconnected pores (average pore size = 49.6 ± 40.9 µm, average window size = 12.6 ± 6.1 µm) and structural characteristics suitable for soft tissue applications. The BJ5ta fibroblast cell line was cultured on scaffolds for 14 days to deposit ECM, after which constructs were decellularized. When tested *in vitro*, periosteal-typical cells exhibited a significantly higher growth rate on ECM-decorated scaffolds compared to controls. Additionally, the chick chorioallantoic membrane (CAM) demonstrated that ECM decoration had a positive angiogenic effect. This proof-of-concept study highlights a promising approach to enhance the biological properties of synthetic membranes, while avoiding challenges associated with decellularizing whole tissues.

## 2. Introduction

The periosteum is the thin, fibrous connective tissue that covers the surface of non-articular bone. It contains a dense network of blood vessels and a diverse cell population comprising stem cells, osteoblasts, and fibroblasts. Studies have shown that the periosteum plays an important role in the natural bone healing process by stabilising fracture fragments [1], facilitating molecular crosstalk between muscle and bone [2, 3], and acting as a source of multipotent progenitor cells [4].

Recently, there has been increasing interest in the development of artificial periosteal membranes that can be used in conjunction with void fillers to facilitate healing in complex bone injuries, such as nonunions and critical-size defects. This interest is partly due to the success of periosteal grafting, where periosteum is harvested from one site and transplanted to a defect site using microvascular anastomoses. Periosteal grafts have been reported to have a success rate of up to 100% in thirteen out of fourteen studies on the treatment of long bone nonunion [5]. However, their widespread adoption has been hindered by several limitations, including graft availability, donor site morbidity, and complexity of surgery [6, 7]. As such, attention has shifted towards the development of an artificial periosteum that can serve as an off-the-shelf alternative, which is a growing area of research [8].

Different strategies for creating artificial periosteal membranes have been explored, several of which involve decellularization techniques. Decellularization is the process of using chemical and mechanical means to remove genetic material from a biological sample while preserving the extracellular matrix (ECM). ECM is the tissue-specific, non-cellular component in all connective tissues, and is composed of highly organised biomolecules [9]. Using decellularization, the aim is to create a periosteal substitute with a familiar microenvironment for cells, while eliminating immunogenic material such as DNA.

Multiple studies have explored methods of decellularizing periosteal tissue [10-13], and it has been demonstrated that it is possible to remove cellular components while retaining ECM and the characteristic double-layered structure of the periosteum [14, 15]. Though not explicitly tested for the purpose of creating an artificial periosteum, this approach could be a promising technique. For example, He et al. showed that a decellularized periosteum prevented fibrous invasion and promoted bone regeneration when used in a rabbit skull defect model [13]. Similarly, Zhang et al. found that decellularized periosteum seeded with mesenchymal stem cells functioned as an osteogenic and angiogenic scaffold, resulting in accumulated vascular endothelial growth factor (VEGF) and a higher local expression of osteogenic markers such as BMP-2 and alkaline phosphatase when implanted in mice [12]. However, this method still requires that large amounts of periosteum tissue be harvested. Moreover, due to its dense collagen network, native periosteum may exhibit low cell permeability and poor regulation of early inflammation [16].

To overcome this, other more readily available decellularized tissues have been tested as periosteal grafts, such as porcine small intestinal submucosa (SIS). SIS is made up of 90% collagen and is known for being elastic, biodegradable, and highly bioactive [17]. It has been tested for its potential as a periosteal graft with conflicting results. Moore et al. found that periosteal sleeves made from laminated and dried SIS could not induce new bone formation in a rat critical size defect model [18], while Su et al. found that combining SIS with PCL via shell-core electrospinning supported both angiogenesis and osteogenesis [19]. These findings suggest that SIS may offer biological potential, though not the required physical properties to act as a periosteal substitute.

As an alternative to decellularizing whole tissues, decoration of more mechanically suitable, synthetic materials with ECM is a strategy that can be used in cases where the base polymer has limited biological properties. The incorporation of selected exogenous ECM components (e.g. peptides, proteins, growth factors) alone is not sufficient to mimic the complexity and unique composition of tissue-specific ECM [20]. As such, decoration with *in vitro* cell-derived ECM via decellularization processes has become an area of increasing interest [21].

In this study, we investigate the use of fibroblast-derived ECM-decorated porous membranes as synthetic periosteal grafts. To the best of our knowledge, this is the first example of *in vitro-generated* ECM being assessed for use in periosteal substitutes. Porous scaffolds were fabricated using an emulsion templating technique with a blend of biodegradable polymers: polycaprolactone methacrylate (PCL-M) and poly(glycerol sebacate) methacrylate (PGS-M). Porous scaffolds were populated with a fibroblast cell line and cultured to allow ECM deposition. Subsequently, scaffolds were decellularized to remove immunogenic material. We assessed the scaffolds’ ability to support attachment and proliferation of cells that are relevant to the periosteum, and evaluated their angiogenic response using an *ex-ovo* chick chorioallantoic membrane assay.

## 3. Materials and Methods

### 3.1. Materials

Photoinitiator (2,4,6-Trimethylbenzoyl Phosphine Oxide/2-Hydroxy-2-Methylpropiophenone blend), resazurin sodium salt, L-glutamine, trypsin-EDTA, penicillin/streptomycin, foetal bovine serum (FBS), DPX mounting media, and haematoxylin solution were purchased from Sigma Aldrich (Poole, UK). Chloroform (99%), toluene (99.5%), ethanol (99%), methanol (99%), phosphate buffered saline (10X solution), fungizone, Quant-iT dsDNA high sensitivity assay kit and Dulbecco’s modified eagle medium (DMEM) were purchased from Fisher Scientific (Pittsburgh, PA). The surfactant, Hypermer B246 (98%), was received as a sample from Croda (Goole, United Kingdom). Eggs were purchased from MedEggs (Norfolk, UK). Eosin solution was purchased from Acros Organics.

### 3.2. Preparation and Characterisation of PCL-M:PGS-M PolyHIPEs

Polymer high internal-phase emulsions (polyHIPEs) were prepared using a 50:50 ratio of PCL-M and PGS-M. PCL-M and PGS-M were synthesised in-house using methods that have previously been reported [22, 23].

PolyHIPEs were made by mixing equal weights of PCL-M and PGS-M (0.5 g total) with Hypermer surfactant (10% w/w surfactant/polymer blend). The mixture was heated until the surfactant was completely melted. The mixture was then cooled to room temperature, and 0.5 g of a 60:40 (w/w) chloroform/toluene solvent mixture was added to reduce the viscosity. The mixture was stirred in a water bath at 50 °C until everything had combined. The emulsion was formed by adding 4 mL of deionised water dropwise (~1 drop/s) with continued stirring. Then, 5% (w/w) photoinitiator was added and the mixture was stirred for a further 5 minutes before being transferred into circular PDMS moulds. The moulds were placed under UV light for 90 seconds to cross-link the polymer. Solid polyHIPE scaffolds were cut to reveal the internal porous structure, washed three times in methanol, once in 70% ethanol, then a further three times in sterile PBS to prepare them for cell culture.

To characterise the porous structure of the polyHIPE scaffolds, scanning electron microscopy (SEM) was used. Samples were dried overnight and gold sputter-coated at 15 kV for 5 minutes. Samples were imaged using an FEI Inspect F SEM with 3 kV power. Three images were taken at three different locations. On each image, 60 pores and 60 windows were measured within ImageJ. Pores and windows were chosen for measurement by overlaying an 8 by 8 grid onto the image, and pores or windows that were in contact with the grid were measured. A statistical correlation factor of 2/ √3 was applied to correct for the underestimation of diameter caused by uneven sectioning [24, 25].

### 3.3. Cell Seeding on PolyHIPE Scaffolds

BJ5ta cells, a male human foreskin fibroblast cell line, were used to produce matrix on the scaffolds. Y201 cells and MLOA5 cells, a human mesenchymal stem cell line [26] and mouse late-stage osteoblast cell line, respectively, were used to assess cell behaviour on decellularized scaffolds. All cells were grown in expansion media (EM) composed of DMEM supplemented with 10% foetal bovine serum (FBS), 2 nm L-glutamine, and 100 mg/mL penicillin/streptomycin. Cells were trypsinised, counted, and resuspended in cell culture media to form a cell suspension with a concentration of 20,000 cells/20 μL. Before cell seeding, polyHIPE scaffolds were soaked in EM for 1 hour in the incubator, then placed into a 24-well plate and partially dried in sterile conditions for 30 minutes. 20 μL of cell suspension was added to each scaffold. Tissue culture plastic (TCP) was used as a positive control. Scaffolds were placed into the incubator for 45 minutes to allow cells to form attachments to the scaffold. Finally, 1 mL of EM was carefully added to each well, and scaffolds were returned to the incubator. This was considered to be day 0, and EM was changed every other day unless otherwise specified. In the case of BJ5ta cells, EM was supplemented with 50 μg/mL ascorbic acid to promote ECM deposition.

### 3.4. Matrix Production on PolyHIPE Scaffolds

Deposition and attachment of collagen onto polyHIPEs was measured using Sirius red staining (SRS), a strong anionic dye that binds to collagen molecules. Cells were cultured for up to 28 days to assess how much collagen was produced, and scaffolds were also tested before and after the decellularization process to assess whether it impacted deposited ECM. EM was removed and scaffolds were washed twice with PBS before being fixed with 3.7% formaldehyde solution for 30 minutes. Scaffolds were washed twice more with PBS, then submerged in 1 mL of SRS solution and left for 18 hours with orbital shaking. The solution was then removed, and scaffolds were washed gently with water until clear. Scaffolds were submerged in a known volume of 0.2 M NaOH/MeOH (1:1) to destain and left for 30 minutes at room temperature with orbital shaking. 150 μL of the destain solution in triplicate was transferred to a clear 96-well plate and the absorbance was read at 405 nm. A fresh scaffold was used at each time point.

### 3.5. Decellularization of PolyHIPE Scaffolds Populated with Fibroblasts

Three different combinations of methods for decellularization were assessed on BJ5ta-seeded polyHIPE scaffolds: freeze-thawing (ft), ft + Triton (t), ft + t + DNase. Before each method, EM was removed and samples were washed twice with PBS. The method of ft can be considered as mechanical decellularization, and works on the principle of intracellular ice crystal formation that assists with cell lysis. Here, three consecutive freeze-thaw cycles (20 minutes at −80 °C, 15 minutes at 37 °C) were applied.

Triton is considered to be a chemical decellularization agent and works by disrupting lipid-lipid and lipid-protein interactions. It is a non-ionic detergent and therefore less damaging to ECM components than ionic detergents. When used, scaffolds were incubated with 1 mL of Triton (0.5%) solution for 10 minutes at 37 °C after freeze-thaw cycles were complete. The solution was removed afterwards.

Finally, DNase is an enzymatic decellularization agent which is often used in combination with complementary methods to break down DNA fragments. In the final condition, scaffolds were incubated with 1 mL of DNase solution (0.2 mg/mL) in an incubator for one hour.

After each of the three methods (ft, ft + t, ft + t + DNase) was completed, scaffolds were washed twice in warm PBS to gently remove the cellular component. All methods were completed under sterile conditions. Resulting scaffolds are referred to as ‘decellularized scaffolds’ from this point.

### 3.6. DNA Quantification

The effectiveness of the decellularization process was assessed by measuring DNA content using a Quant-iT dsDNA high sensitivity assay kit. Samples were washed twice with PBS, then 1 mL of cell digestion buffer (10 mM Tris-HCl, 1 mM ZnCl2 and 1% Triton X-100 in distilled water) was added and incubated at room temperature for 1 hour. Samples were vortexed for 60 s and kept in the fridge overnight at 4 °C. The next day, samples underwent three freeze-thaw cycles (20 minutes at −80 °C, 15 minutes at 37 °C, 15 s vortex) to create the final lysate solution. 10 μL of lysate was added to 90 μL of assay working solution in triplicate in black 96-well plates. The well plate was shaken for 10 seconds to ensure thorough mixing, then incubated for 10 minutes at room temperature to allow the DNA to conjugate fully. A plate reader was used to read fluorescence with an excitation wavelength of 485 nm and an emission wavelength of 535 nm.

### 3.7. Cell Viability and Proliferation of MSCs and Osteoblasts on Decellularized Scaffolds

Decellularized scaffolds were seeded with MLOA5 cells and Y201 cells to assess biological behaviour. Resazurin reduction (RR) assay was used to measure metabolic activity of cells seeded on the scaffolds as a way of estimating cell viability and proliferation. The RR assay works on the mechanism in which the blue, nonfluorescent salt resazurin gets reduced by live cells and forms pink, fluorescent resorufin, which is detectable by a fluorescence plate reader. EM was removed from the wells, and 1 mL of resazurin solution was added to the scaffolds in dark conditions. Well plates were kept for 4 hours in a cell culture incubator. From each well, 150 μL of the reduced solution in triplicate was transferred to a 96-well plate. The fluorescence was measured in the plate reader at an excitation wavelength of 540 nm and an emission wavelength of 590 nm. The RR assay was performed on days 1, 4, and 7. A fresh scaffold was used for each time point to negate the effects of resazurin exposure on cells.

### 3.8. *Ex-Ovo* CAM Assay

The *ex-ovo* chicken chorioallantoic membrane (CAM) assay was used to assess cytotoxicity, cell infiltration, and angiogenic response of decellularized scaffolds. Briefly, pathogen-free fertilised eggs (Shaver Brown, MedEggs) were cleaned and incubated at 37 °C in a humidified hatching incubator for 3 days. On day 3, eggs were carefully cracked into sterile weighing boats containing 3 mL of PBS + 1% (v/v) penicillin streptomycin solution and 1% (v/v) fungizone. Eggs were then kept in a cell culture incubator. On day 7, decellularized scaffolds were placed onto the surface of the CAM region of the eggs. Non-decellularized polyHIPE scaffolds (cell and ECM-free) were used as a control. Eggs were incubated for a further 7 days. On day 14, images were taken of the sample and surrounding vasculature. All embryos were then sacrificed via beheading. Scaffolds and surrounding membrane tissue were excised for histological analysis.

### 3.9. Haematoxylin and Eosin (H&E) Staining

Samples taken from the CAM assay were washed three times in PBS and then fixed with 3.7% formaldehyde solution for 24 hours. The samples were again washed with PBS, then placed into tissue cassettes and processed for 18 hours (Leica TP 1020 tissue processor, Leica Biosystems, UK). The samples were embedded in wax, and 10 μm-thick sections were taken. Sections were stained using haematoxylin for 90 seconds and eosin for 3 minutes, washed with tap water, dehydrated via a series of washes in industrial methylated spirits (IMS) before finally being washed twice with xylene and mounted with DPX. The sections were imaged using a light microscope.

### 3.10. Haematoxylin and Eosin (H&E) Staining

Statistical analysis was completed using statistical analysis software (GraphPad Prism, Version 8.4.3, CA, United States). All datasets were analysed using a one-way or two-way analysis of variance (ANOVA) followed by a Tukey multiple comparison test. Error bars on graphs indicate standard deviation, and the number of technical repeats (n) is given in figure captions where relevant. Number of experimental repeats (N) was 2 for all biological experiments.

## 4. Results and Discussion

### 4.1. Fabrication of Decellularized PolyHIPE Scaffolds

PolyHIPE scaffolds made from a blend of PCL-M:PGS-M were chosen due to their mechanical flexibility and ease of synthesis. The average pore size and window diameter were measured to be 49.6 ± 40.9 µm and 12.6 ± 6.1 µm, respectively, though a large spread of pore sizes was observed. The formulation of the emulsion was previously optimised by our group to maximise pore size, window size, and window frequency by varying parameters such as the temperature of the external phase during emulsion formation and the solvent blend used. As such, the polyHIPE not only has large pores but also large and frequent interconnects between the pores (Figure 1A and 1B), making it ideal for supporting cell infiltration and nutrient transfusion.

**Figure 1:**
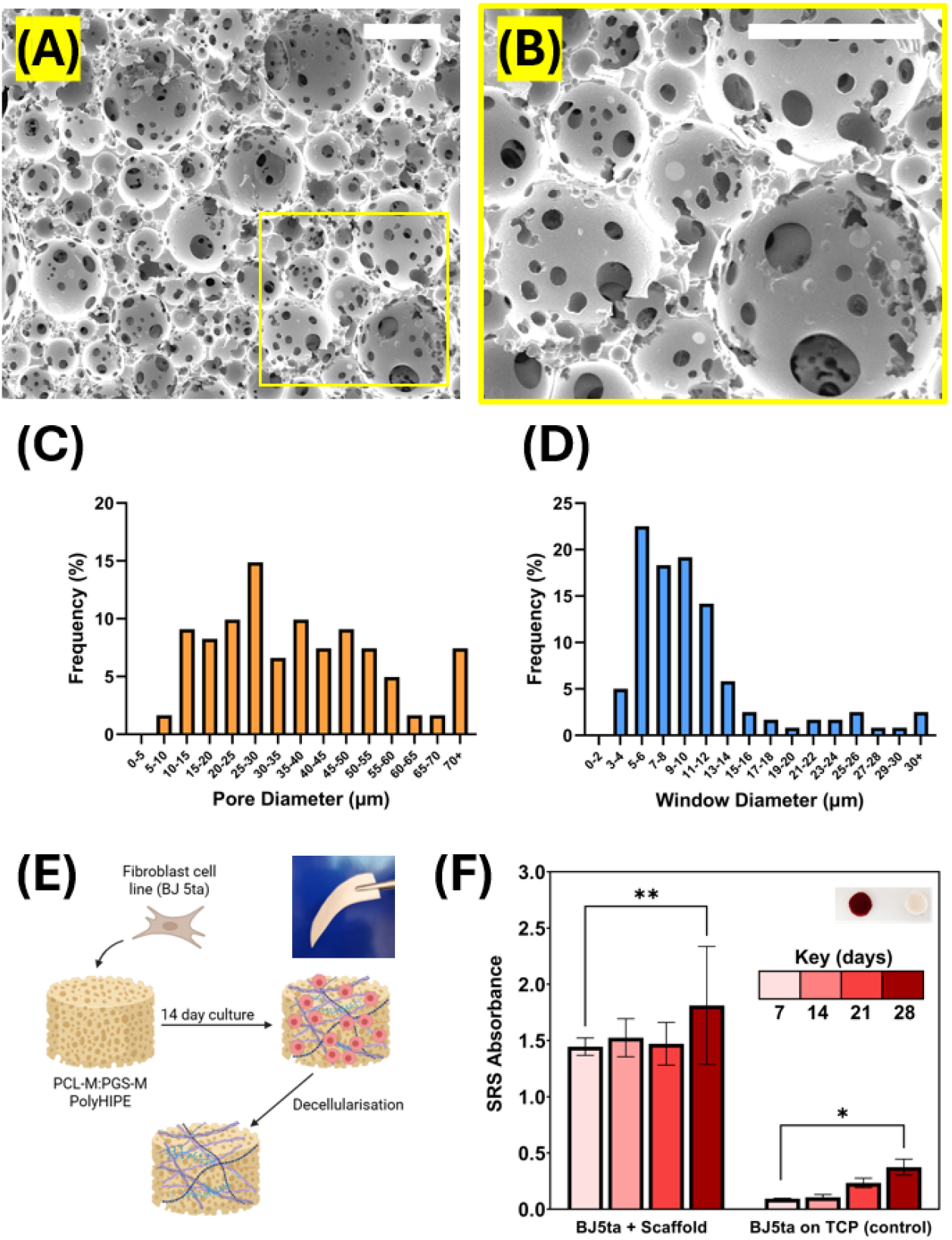
(A) SEM micrograph of PCL-M:PGS-M polyHIPE internal structure with (B) highlighting the interconnecting windows between pores. Scale bars represent 150 µm. Graphs show the distribution of (C) pore sizes within the structure and (D) the window sizes. (E) The process for creating decellularized polyHIPE scaffolds. Upper right picture shows the flexibility of the polyHIPE membrane. (F) Collagen deposition by BJ5ta cells on polyHIPE scaffolds over time as demonstrated by SRS. Upper right shows SRS-stained cell-seeded scaffolds on day 28 vs cell-free control (n=3, ^*^P < 0.033, ^**^P < 0.002).

BJ5ta cells were chosen because fibroblasts are one of the major cell populations found within the periosteum, and these are a commercially available human fibroblast cell line that can produce ECM [27]. Fibroblasts produce and secrete all components of the ECM, including structural proteins, adhesive proteins, and space-filling ground substance that is primarily composed of glycosaminoglycans and proteoglycans [28]. Collagen production by BJ5ta cells on polyHIPE scaffolds was assessed via SRS (Figure 1F) and was observed to be rapid, with no significant difference between time points occurring until day 28. SRS also demonstrated that collagen deposition was consistent across the surface of the scaffold, likely due to the fast-growing nature of the cells and the relatively high cell seeding density used. The amount of collagen produced by day 14 was deemed suitable, and this time point was therefore used to decrease overall production time of decellularized scaffolds.

### 4.2. Decellularization of PCL-M:PGS-M PolyHIPE Scaffolds Seeded with Fibroblasts

Different decellularization techniques were tested to reduce remaining DNA content to the lowest value possible, therefore reducing the risk of immunogenic response from the host once implanted. There are several methods for decellularizing tissues as reported in literature, which generally work by destroying the cell membrane in order to remove cellular material [29]. Whole organs and tissues may undergo rigorous decellularize processes, but for *in vitro*-generated ECM, more gentle methods are used in order to better preserve deposited ECM on the scaffolds [30].

The efficiency of ft, ft + t, and ft + t + DNase as decellularization treatments was tested, and the remaining DNA amounts were measured at 65.5%, 45.4%, and 11.4% of the total amount of initial DNA, respectively (Figure 2A). A significant difference in DNA amount was observed in the ft + t + DNase group; hence, this was chosen for its ability to remove up to 89% of DNA content. The gold standard set by Crapo et al. in 2011 states that tissues are considered to be decellularized once dsDNA content is < 50 µg per mg of dry tissue [31]. With this group, remaining DNA content was measured to be less than 50 µg across the entire scaffold (average dry weight = 70.6 mg) hence the decellularization method was deemed appropriate.

**Figure 2:**
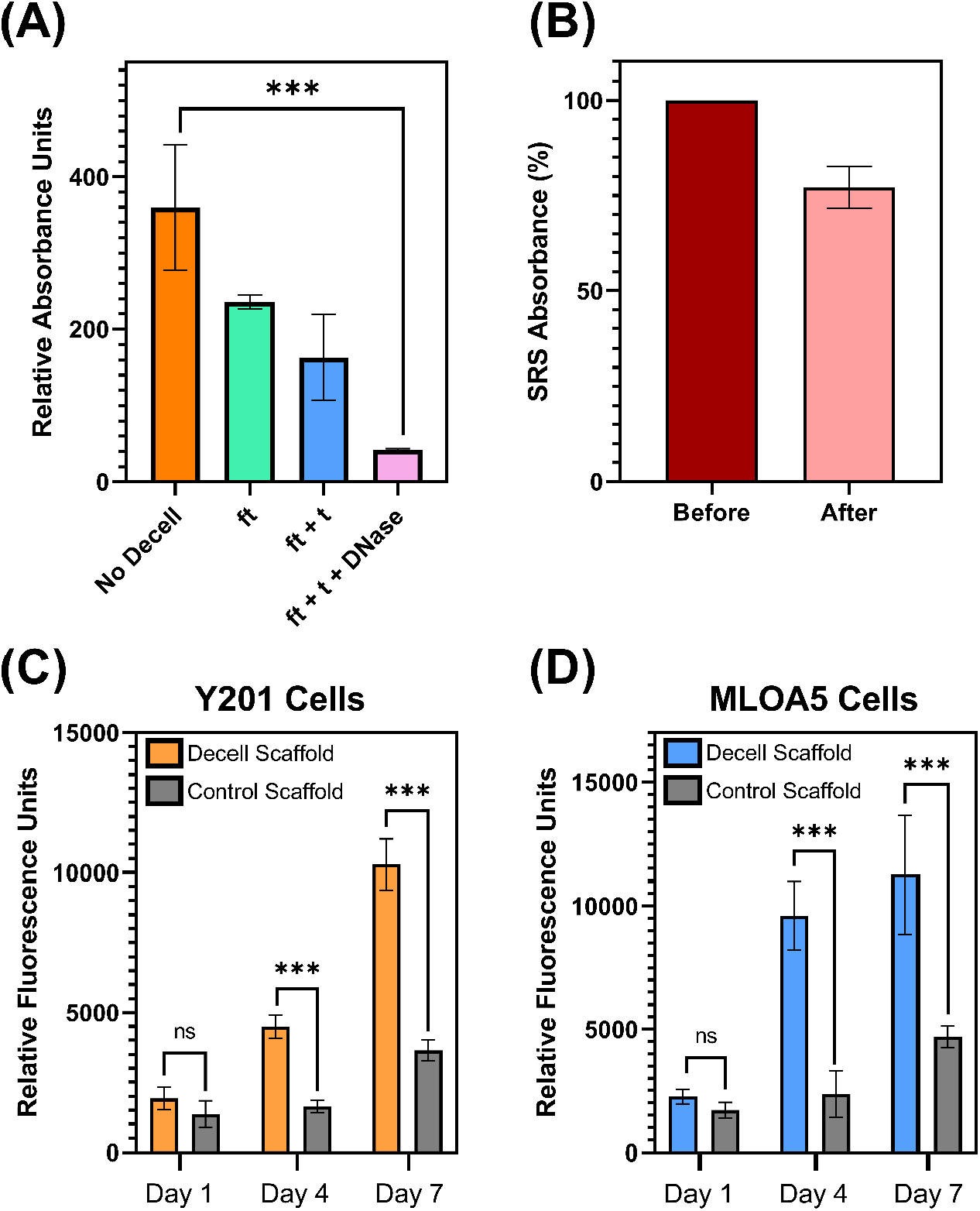
(A) Comparison of various decellularization techniques as shown by remaining DNA content. (B) Collagen content of polyHIPE scaffolds seeded with BJ5ta cells for 14 days before and after decellularization. Metabolic activity of (C) Y201 cells and (D) MLOA5 cells on decellularized polyHIPE scaffolds. (n = 3, ^***^P < 0.001, ns = no significance).

A similar method was tested by Dikici et al. for decellularization of 3D-printed PCL-M polyHIPE scaffolds with an osteoblast-like matrix [21]. They found that ft + DNase only was able to remove up to 95% of DNA content without the need for Triton, a higher rate than observed in this study. This could be due to the larger macropores within their structure making it easier for decellularization agents to reach cells, and for washing steps to remove cellular components. One option to improve decellularization within this study could be to use further washing steps to ensure complete removal of released cellular components.

The decellularization process involves several steps, each with washing stages in between. As such, there are several opportunities for deposited ECM on the polyHIPE scaffolds to be disturbed or removed entirely. Following decellularization via ft + t + DNase, it was found that 77% of collagen was preserved on polyHIPE scaffolds, with no significant difference found between the amount of collagen before and after decellularization.

### 4.3. *In vitro* Response of Cells to Decellularized PolyHIPE Scaffolds

In separate cultures, Y201 cells and MLOA5 cells were seeded onto decellularized polyHIPE scaffolds to assess their response in comparison to polyHIPE scaffolds with no ECM components. These cells are representative of two major cell populations within the periosteum that play an active role in bone healing: mesenchymal stem cells and osteoblasts, respectively. Metabolic activity of the cells was measured by the resazurin reduction assay (Figure 2C and 2D), which was used to assess cell attachment, viability, and proliferation over time.

Day 1 results from the resazurin reduction assay can be used as an indication of attachment, when cells have undergone a maximum of one round of cell doubling. Decoration of materials with ECM is thought to increase cell attachment due to increased cell adhesion sites and increased surface roughness [32, 33]. For both Y201 cells and MLOA5 cells, there was no significant difference in measured metabolic activity after 24 hours when comparing decellularized versus controls, showing equal seeding efficiencies. This is likely because the polyHIPE blend already has very good cell attachment properties, with the addition of PGS-M to the polymer blend increasing hydrophilicity and the number of cell adhesion sites [34].

On days 4 and 7, metabolic activity from both cell types was significantly higher on decellularized scaffolds compared to controls, with day 7 showing a nearly three-fold improvement. This provides further evidence that polyHIPE scaffolds retain important ECM components following the decellularization process. The presence of ECM proteins on material surfaces has previously been demonstrated to have a positive impact on cell growth [35], regardless of whether the ECM-depositing source matches the seeded cell type [21]. ECM is specific to tissues, with each cell type generating unique forms of ECM, but common components of ECM, such as collagens and adhesive glycoproteins, are beneficial to many cell types [36]. Additionally, growth factors such as platelet-derived growth factor (PDGF) and transforming growth factor beta (TGF-β) can bind to components of ECM [37]. The gentler decellularization processes used on the *in vitro*-generated ECM may permit retention and functionality of these growth factors, improving bioactivity.

### 4.4. *In vivo* Response of Cells to Decellularized PolyHIPE Scaffolds

The *ex-ovo* CAM assay is a simple and cost-effective *in vivo* model that can be used to screen for beneficial tissue behaviours in response to a material including cell infiltration, cytotoxicity, and vascularisation [38]. It is reported that the average embryo survival rate for intermediate users is 68%, hence this should be the comparison for whether a material is potentially toxic [39]. In this study, the embryo survival rates for decellularized scaffolds and polyHIPE-only scaffolds were 75% and 81% respectively, demonstrating that there was no adverse effect on embryo survival caused by the material.

At the stage of embryonic development at which the CAM assay is completed, the chicken embryos are considered to be immunodeficient [40]. This means that the model can not be used to assess whether decellularized scaffolds have a low enough genetic content to prevent immune response. It is, however, a good indication of whether scaffolds remain sterile through the decellularization process, as CAM embryos may become infected if they come into contact with non-sterile materials. In this case, embryos that died did not have infection present around the scaffold at the time of sacrifice, hence scaffolds were assumed to be sterile.

Angiogenesis is an essential part of the natural healing process, and within bone tissue engineering as a whole, there is increasing interest in stimulating angiogenesis at the defect site. PCL-M:PGS-M polyHIPE scaffolds support cell ingrowth but are not angiogenic; hence were used as a negative control to allow for observation of the angiogenic impact of deposited ECM. By Day 14, all polyHIPE samples were well integrated into the CAM and had to be cut away from the membrane during isolation. Decellularized samples were occasionally partially enveloped by the CAM, as seen in Figure 3, though this was not observed in the control group. Directional vasculature growing towards implanted decellularized scaffolds was observed, whereas the vasculature surrounding polyHIPE-only controls appeared randomly orientated. Histological analysis showed cellular infiltration on both polyHIPE-only scaffolds and ECM-decorated polyHIPE scaffolds, though more infiltration was present on the latter (Figure 3). Additionally, small blood vessels could be seen growing into ECM-decorated scaffolds.

**Figure 3:**
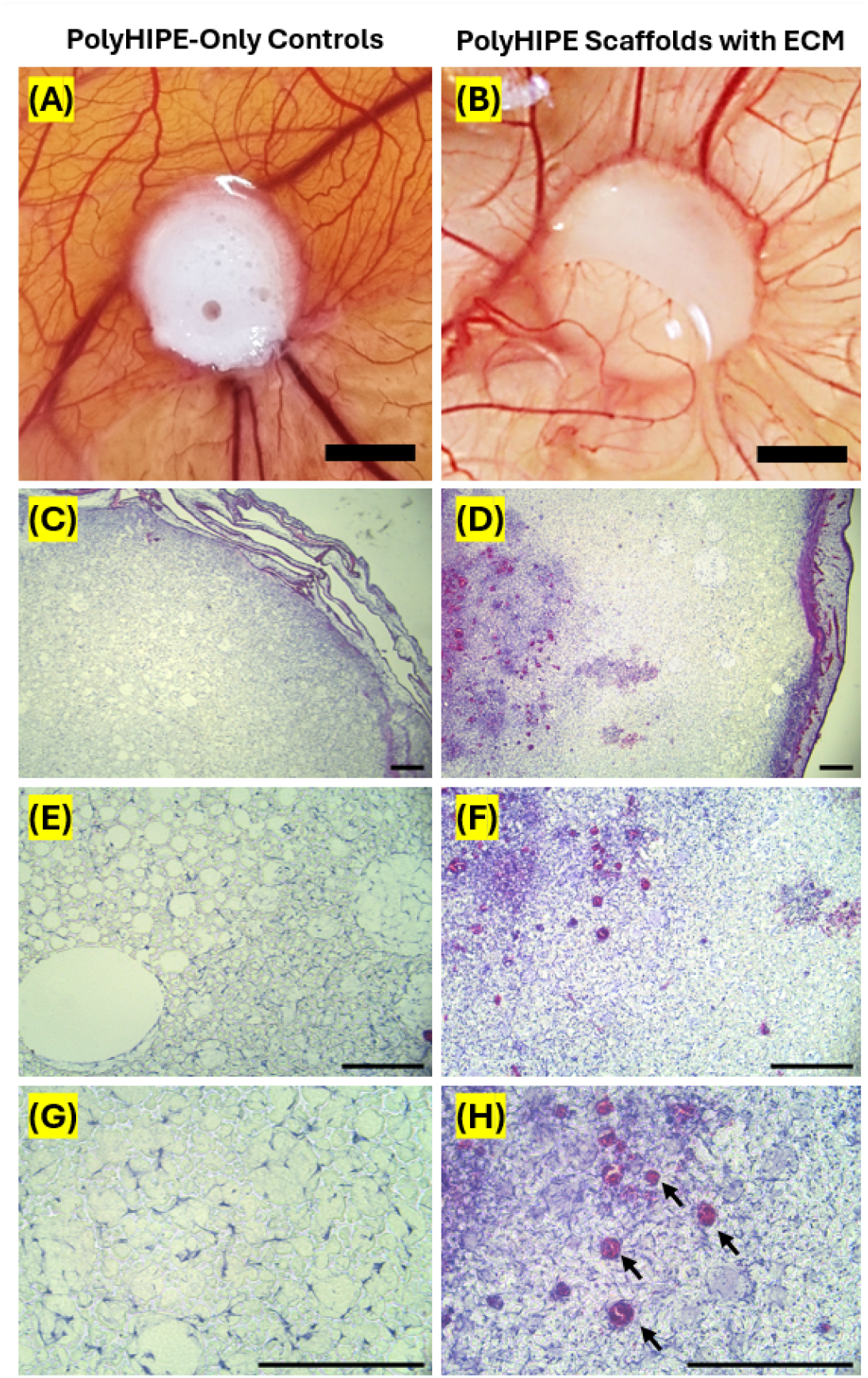
Angiogenic assessment of PCL-M:PGS-M-only polyHIPE scaffolds vs polyHIPE scaffolds containing ECM using *ex-ovo* CAM assay (A and B). Scale bars represent 500 µm. Images were taken on day 14 of embryonic development. Histological analysis of scaffolds removed after CAM assay at (C and D) 4✕, (E and F) 10✕, and (G and H) 20✕. Arrows point to blood vessels. Scale bars represent 250 µm.

It has previously been shown that *in vitro-generated* ECM has a positive angiogenic effect when tested *in vivo*. Pham et al. demonstrated that increasing amounts of ECM generated *in vitro* on titanium fibre mesh scaffolds by MSCs resulted in increased vascular invasion when implanted in a rat animal model [30]. This was thought to be due to angiogenic factors released from the ECM, which may include VEGF and BMP-2. Some VEGF isoforms can bind to ECM and may induce angiogenesis either whilst still bound to the ECM or once processed by proteases [41], further supporting this argument.

## 5. Conclusion

This study demonstrates the potential of using in vitro-generated, fibroblast-derived ECM to improve the biological functionality of polyHIPE membranes to act as synthetic periosteal grafts. By decorating porous PCL-M:PGS-M scaffolds with cell-derived ECM, we created membranes that supported increased proliferation of periosteal-typical cells, resulted in increased cellular infiltration, and exhibited pro-angiogenic behaviour in the CAM assay. These findings suggest that ECM decoration may provide a relatively straightforward and reproducible approach to improve synthetic scaffold bioactivity, without the limitations associated with decellularizing whole tissues. While this work serves as a proof-of-concept, further studies are required to assess the in vivo performance of these constructs as well as their osteogenic capabilities.

## 6. Acknowledgments

The authors would like to thank the EPSRC-funded Centre for Doctoral Training in Advanced Biomedical Materials grant reference EP/S022201/1 provided to Caitlin Ryan. Electron microscopy was performed in the Sorby Centre for Electron Microscopy at the University of Sheffield

